# Direct observation of secondary nucleation in huntingtin amyloid formation by High-Speed Atomic Force Microscopy

**DOI:** 10.1101/2024.11.05.622155

**Authors:** C. van Ewijk, G. Jain, Y. K. Knelissen, S. Maity, P.C.A. van der Wel, W.H. Roos

## Abstract

Amyloid fibril formation is a hallmark of various neurodegenerative diseases such as Huntington’s (HD), Alzheimer’s and Parkinson’s disease. The protein aggregation process involves slow nucleation events followed by rapid growth and elongation of formed fibrils. Understanding the pathways of amyloid formation is key to development of novel therapeutic agents that can interfere with the pathogenic protein misfolding events. Recent studies of aggregation by polypeptides from Alzheimer’s and Huntington’s disease have identified the importance of a poorly understood secondary nucleation process that may even be the dominant source of protein aggregate formation. Here we focus on the polyglutamine-expansion disorder HD and employ mechanistic and structural studies to study different aspects of secondary nucleation in the aggregation of huntingtin Exon 1 (HttEx1). Notably, we apply high-speed atomic force microscopy (HS-AFM) to directly observe the process on the single-particle level and in real time. Our observations show unique features of the amyloid formation dynamics in real time, including secondary nucleation, elongation and the formation of large bundles of fibrils as a result of nucleated branching. We examine the role of HttEx1 flanking segments during the aggregation process, revealing that the N-terminal Htt^NT^ segment exhibits a clear primary nucleation-aggregation-enhancing ability, however, that it does not seem to induce or affect the secondary nucleation process. The obtained results illuminate the complex aggregation process of HttEx1 and have implications for attempts to inhibit or modulate it for therapeutic purposes.

## Introduction

Aggregation of peptides and proteins into insoluble amyloid fibrils is a hallmark of various neurodegenerative diseases ^1^. Given the potential pathogenic role of these aggregates, there is a broad interest in understanding the molecular mechanisms of amyloid formation. Amyloid formation is typically considered to be a nucleation-based process that features two distinct phases: the initial phase where (primary) nucleation occurs, and the elongation phase where a rapid growth of existing fibrils can be observed ^2,3^. The elongation process traditionally was explained by elongation of existing fibril ends. It has been seen as a molecular rationale for prion-like propagation, where existing fibrils act as catalysts for subverting additional protein monomers into a pathogenic state. Recently, quantitative analysis of fibril growth kinetics has shown that existing fibrils serve as a nucleation point for new fibrils, in a way that is not explained by fibril elongation alone ^4–7^. This process, called secondary nucleation, has been shown to even be a dominant aggregation mechanism for several different amyloid proteins implicated in disease ^4–6,8,9^. Accordingly, there has been an impetus toward preventing amyloid formation through the inhibition of secondary nucleation ^10–12^. However, for such an effort, a better understanding of the mechanistic details of secondary nucleation is vital, especially for the design of effective therapeutic agents.

Despite its important role, the secondary nucleation process remains poorly understood and is highly debated ^2,13^. To elucidate this process, we turn towards huntingtin (Htt) protein aggregates associated with Huntington’s disease (HD) ^14^. This neurological disease results from a heritable mutation that causes an expansion of a glutamine repeat region (polyQ) in the first exon of the huntingtin protein, leading to intraneuronal protein deposits ^15^. These deposits have been implicated in determining disease onset, progression and toxicity ^16–18^, although the precise impact is thought to depend on the nature of the inclusions ^16^. The cellular inclusions (in patients and model animals) contain N-terminal fragments of the Htt protein, which match the first exon (HttEx1)^15^. The generation of this HttEx1 fragment is driven by aberrant splicing or proteolytic events, which are both enhanced by the repeat expansion of the mutated gene ^19–21^. Consequently, HttEx1 generation increases progressively as a consequence of somatic expansion of the CAG repeat ^22^. HttEx1 features the (mutated) polyQ stretch, flanked by a 17-residue N-terminal segment (Htt^NT^) and a proline-rich C-terminal domain (PRD) (see Fig. 1). These flanking segments impact on the amyloid formation process, and form a disordered fuzzy coat in the aggregates ^23–28^.

**Figure 1:**
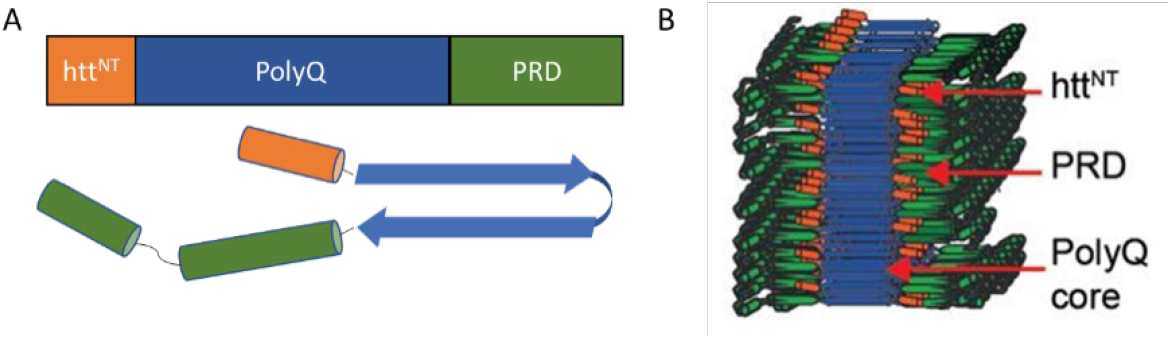
Huntingtin exon 1. A) Primary and secondary structure of huntingtin exon 1 showing the N-terminal domain (Htt^NT^), the polyglutamine core (PolyQ) and the proline-rich domain at the C-terminal (PRD). B) Model of HttEx1 amyloid fibril structure.

The exact role and effect of the flanking segments on the amyloid formation process remains however poorly understood. The Htt^NT^ segment is believed to play a vital role in primary nucleation, bringing the polyQ stretches in close proximity allowing for the misfolding into the β-sheet structure ^26,29^. However, there are multiple conceptions on the effect of the N-terminal domain on the aggregation propensity and morphology of HttEx1 fibrils. It has been reported to enhance HttEx1 aggregation ^27,30,31^, an effect which is even increased upon acetylation of Htt^NT 32^. Studies where Htt^NT^ was removed report varied effects, from a lack of bundled fibrils ^31,33,34^, to a relative increase of lateral association and clumping ^26^. It has also been suggested that parallel Htt^NT^ on the side of an existing fibril are the driving factors for secondary nucleation and subsequent branching ^29^. The reported polymorphism of HttEx1 fibrils is attributed to the fuzzy coat of the flanking domains and to local environmental conditions ^33–36^. Such polymorphism results in different levels of neurotoxicity, although a mechanistic explanation is still lacking ^37,38^. Speculations on the role of secondary nucleation on the emergence of polymorphism and toxicity of the deposits created high interest in understanding the underlying molecular processes to allow for possible control and inhibition of various aspects of amyloidogenic disorders which are currently incurable.

The aggregation behavior of HttEx1 shares common features with other amyloidogenic proteins, as β-sheet-rich nuclei are formed after a lag phase followed by a rapid increase of fibril lengths ^27,33,37^. By introducing preformed fibrils the slow lag phase can be bypassed, a ‘seeding’ process which has been implicated to play a vital role in the disease progression of amyloid-related neurodegenerative disorders ^39,40^. Transmission of β-sheet-rich protein deposits between neurons has been implied in disease progression of HD patients ^39,41^. Unlike, for example, amyloid Beta 42, huntingtin amyloids form bundles of fibrils with a branched structure as observed by atomic force microscopy (AFM) ^7,42^ and electron microscopy (EM) ^33,36^. It has been speculated that the bundles of HttEx1 fibrils are not formed due to lateral association of existing filaments, but may be a result of secondary nucleation along a primary fibril surface ^7^, which implies that secondary nucleation plays a crucial role in the polymorphism of HttEx1 fibrils. Combined, these features make HttEx1 an appealing system to study the elusive secondary nucleation process in amyloid formation.

To elucidate the huntingtin amyloid formation process we turn towards High-Speed Atomic Force Microscopy (HS-AFM), a single particle technique that allows for label-free, real time observation of dynamic molecular processes with sub-second temporal resolution ^43–45^. HS-AFM has been deployed before to study the formation of other amyloid fibrils, e.g. those implicated in Alzheimer’s disease ^46–48^. Here, we use HS-AFM to capture HttEx1 amyloid formation, thereby unraveling elongation and secondary nucleation in amyloid formation in real time and exploring the role of Htt^NT^ in these processes.

## Results

### Secondary nucleation drives HttEx1 aggregation into fibrils

To study the mechanistic aspects of HttEx1 aggregation in vitro, kinetic studies were performed. Amyloidogenic HttEx1 was produced in an aggregation-resistant state as a fusion protein, using maltose binding protein (MBP) as a solubility tag. In this setup, the aggregation can be triggered on demand by the addition of factor Xa (FXa) protease, which releases HttEx1 to allow it to aggregate into amyloid fibrils (see Fig. 1A-B) ^33,49^. In prior work, we established that Q44-HttEx1 aggregation follows a misfolding pathway dominated by secondary nucleation ^33^. A similar fluorometric thioflavin T (ThT) assay of varying concentrations (20μM to 60μM) of Q32-HttEx1 protein was used to analyse its molecular mechanism of aggregation (see Fig. S1). A clear concentration dependence is visible, with variations in the lag phase time, half-times and slopes. Plotting the half-time versus the log of the monomer concentration gives a linear trend (see Fig. S1B). A linear slope, as observed, indicates that the dominant aggregation mechanism doesn’t change with monomer concentration. The processes that could occur are primary nucleation, secondary nucleation, and fragmentation. Fitting the data to these potential mechanisms (see Fig. S1A) shows that the dominant mechanism for the aggregation of Q32-HttEx1 is secondary nucleation ^50^. The global fit to the ThT curves with a single model (secondary nucleation-dominated mechanism) is shown in Fig. S1A. In the case of nucleation-limited aggregation mechanisms, the aggregation process should be accelerated by adding fibril seeds. Indeed, a significant reduction in the lag phase and acceleration in fibril formation was observed upon the addition of preformed fibrils (see Fig. S2A). Thus, HttEx1 aggregation is a prime example of an amyloid formation process in which there is a central role for secondary nucleation, which is in line with prior studies ^7,33^.

### Morphology of HttEx1 aggregates by TEM and AFM

To have a better understanding of the structural transformations of HttEx1 during the aggregation process, monitored by ThT above, we analysed different time points by negative stain transmission electron microscopy (TEM). Samples from an aggregation experiment with 51μM Q32-HttEx1 were taken and frozen in liquid nitrogen. TEM analysis showed the gradual growth of typical HttEx1 fibrils over time (see Fig. S1C). Small fibrillar and globular species were observed in the lag phase of the aggregation process (2 hrs), with the latter being dominant. The globular species were heterogeneous with a varying diameter of 12-30nm (see Fig. S2D), similar to oligomers seen in previous studies ^51,52^. The fibril widths, seen as indicators of HttEx1 fibril polymorphism ^33,53^, were 6 ± 2 nm (n= 71) (see Fig. S1 C-D). By 20 hours, the fibrils become the dominant species, consistent with the ThT assay results (SI Fig. S1C-D). The width of these fibrils was 11± 2 nm (n=104). Over time, the appearance of the growing fibrils changed, with the width approaching 9 ± 3 nm (n= 222) at 71 hours. At later stages, the fibril width did not vary, but their length increased with time. The mature Q32-HttEx1 fibrils after 96 hours had fibrils widths of 8 ± 2 nm (n=166). The mature fibrils that formed at the end of the aggregation process appeared qualitatively similar for Q32- and Q44-HttEx1 (see Fig. S1C-D; S2), except for differences in width expected for changes in the polyQ segment length. Note that the width observed in the negative stain TEM analysis is thought to represent the dimensions of the fibril core (formed by the polyQ segment alone), as the stain cannot penetrate it ^33,54^.

Thus, TEM can provide good-resolution images of the fibril morphology, but it is less suitable for probing the surface characteristics of the fibrils. To allow more detailed visualization of the fibril surface, mature Q44-HttEx1 fibrils were characterized by AFM (see Fig. 2A-B, S3). The average height of 11 ± 2 nm (n=65) of Q44-HttEx1 fibrils as measured by HS-AFM (See Fig. 2D-E) compares well to results from TEM analysis of the same batch of fibrils (6 ± 2 nm, n= 154; Fig. 2 C, F), considering the latter is thought not to include the flanking segments ^54^. Various aggregate-like structures are visible along the length of the fibrils (see Fig. 2B), indicating that the surface of the fibrils is prone to associate with free monomers or small oligomers. It is worth noting that HttEx1 fibrils are well known to be more heterogeneous and less smooth than other amyloid fibrils, e.g. shown by AFM, cryo-EM and cryo-electron tomography ^7,34,36^.

**Figure 2:**
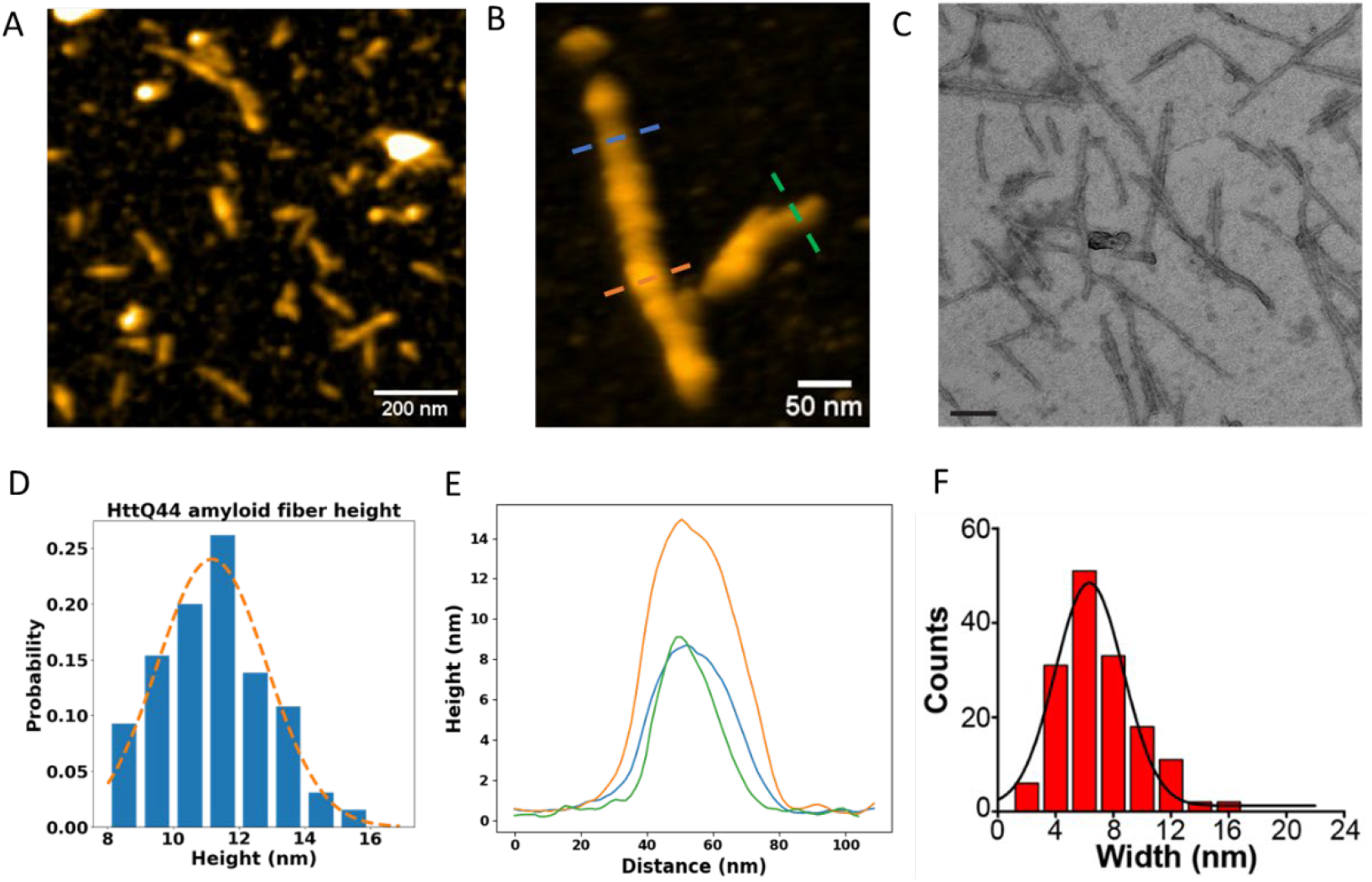
Q44-HttEx1 amyloid fibril morphology by HS-AFM and EM. A) HS-AFM overview image of 60 μM Q44-HttEx1 fibrils on a mica surface. B) Close up of Q44-HttEx1 amyloid fibrils. C) Negative stain TEM image of 60 μM Q44-HttEx1 fibrils formed at room temperature. Scale bar 100nm. D) Distribution of Q44-HttEx1 amyloid fibril height by HS-AFM with an average of 11 ± 2 nm E) Height profiles taken along the drawn lines in (B). F) Distribution of Q44-HttEx1 fibril widths as measured by TEM 6 ± 2 nm.

### Observing secondary nucleation by AFM

Static AFM images could not answer the question of how these characteristic fibril structures form during HttEx1 aggregation. To probe this, the seeded aggregation process of Q44-HttEx1 (see Fig. S2) was followed over time using HS-AFM, upon addition of a solution containing freshly cleaved protein monomers. The evolution of a fibril seed over time was observed by HS-AFM, as displayed in Fig. 3A-B. The seeds elongate bi-directionally with an average rate of 5 ± 3 nm min^-1^ (see Fig. 3C), although occasionally one side of a fiber did not elongate at all. Fibril elongation was generally not continuous over the course of the measurements, with phases of rapid growth alternated by stagnation phases (see Fig. 3B).

**Figure 3:**
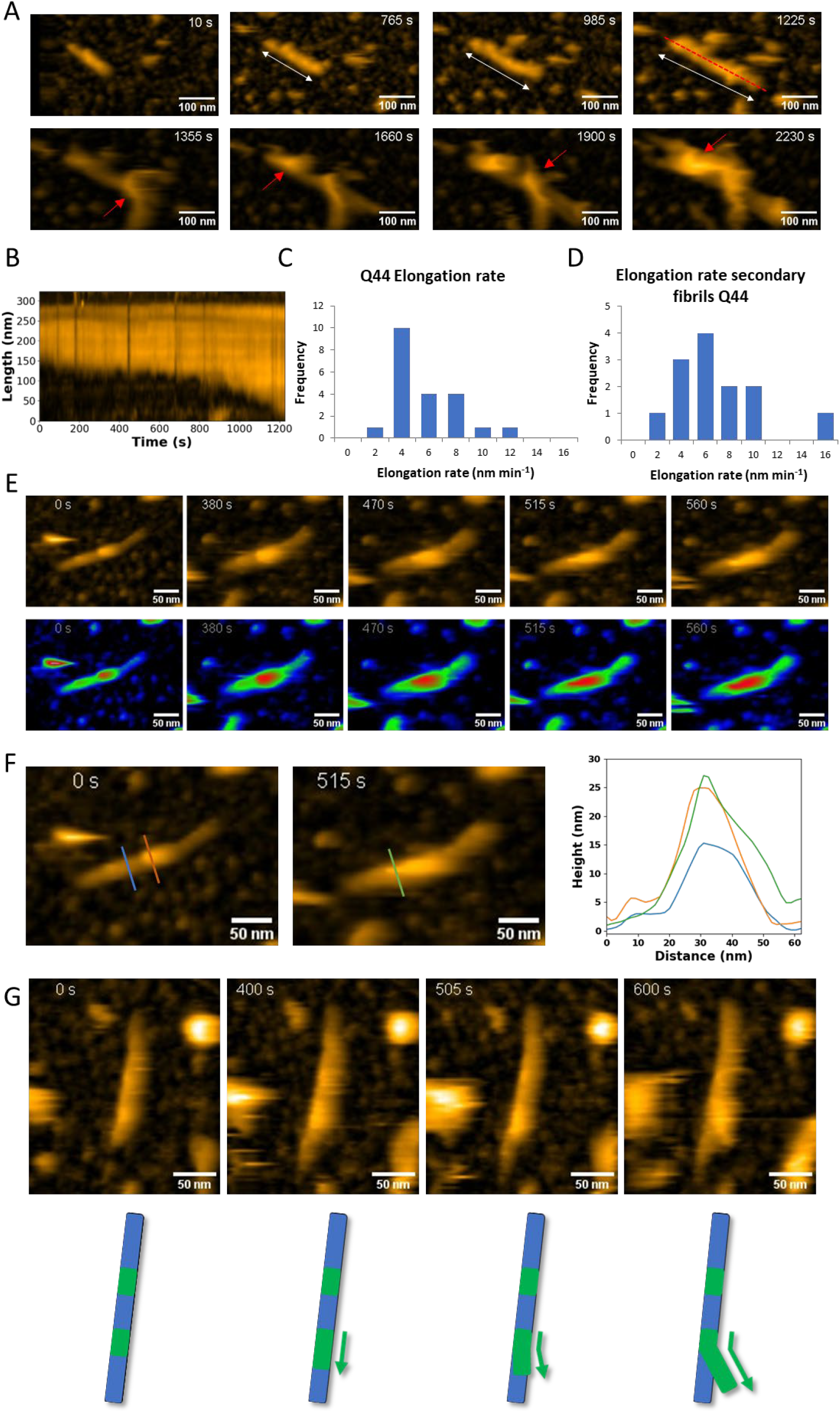
Seeded huntingtin amyloid formation in real time by HS-AFM. A) Snapshots of Q44-HttEx1 amyloid formation showing elongation and nucleated branching. White arrows indicate the elongation of the fibril and red arrows indicate the appearance of secondary nucleated fibrils. Z-scale 0-1225s 35nm, 1355-2230s 80nm. B) Kymograph along the dashed red line in A) showing fibril elongation. C) Elongation rates of Q44-HttEx1 fibrils. Average: 5 ± 3 nm min^-1^ (n=21). D) Elongation rates of secondary Q44-HttEx1 fibrils. Average: 6 ± 3 nm min^-1^ .(n=13). E) Secondary nucleation event on the surface of an Q44-HttEx1 amyloid fibril. In the lower panels, the parent fibril and secondary fibril are coloured in green and red respectively. F) Cross sections of the growing secondary fibril. F) Snapshots of branching of secondary fibrils and corresponding schematic. The green segments indicate the secondary fibrils.

Unlike bulk ThT assays, HS-AFM results allow the observation of molecular events underpinning the overall aggregation kinetics. One striking observation is the detection of secondary nucleation events on the fibril surface, followed by the growth of new fibrils (see Fig. 3E-F, S4). Initially, small aggregates are observed on the fibril surface, manifesting as a sudden increase in the local fibril height (Fig. 3E). Over time, these new features are seen to elongate, adopting a morphology that resembles a fibril structure, with the height of the secondary fibrils similar to that of singular fibrils (see Fig. 3F). Just like primary fibril growth, secondary nucleation also occurs bi-directionally. Furthermore, elongation of these newly formed fibrils occurs along the surface of the parent fibril. The rate of elongation has an average value of 6 ± 3 nm min^-1^ (see Fig. 3D), which is similar to the rate of elongation of the original seed fibrils (above). Eventually, the growing end of the new fibril detaches from the fibril surface, resulting in a branched structure (see Fig. 3G). Multiple secondary nucleation events have been observed along a single fibril to finally appear as a unordered broomlike structure, with the fibrils sticking out at the end of the cluster (see Fig. S5). As noted above, this type of morphology seems characteristic of HttEx1 fibrils. The fact that multiple fibrils originate from a single seed correlates well with the exponential growth observed in the ThT assays, and provides a molecular rationale for the secondary nucleation process identified by kinetic analysis. Regarding the overall prevalence, secondary nucleation occurred on almost all of the fibril seeds over time (∼90%), indicating the high susceptibility of the fibril surfaces to catalyse fibril nucleation events. In an attempt to quantify the amount of amyloid material, we measured the relative volumetric increase over time, which showed an exponential trend corresponding to the shape of the kinetics curve (ThT assays) (see Fig. S6).

No primary nucleation events were observed during the timescales at which elongation and secondary nucleation were captured. These findings are consistent with the bulk aggregation mechanism (see Fig. S1A), which identified a dominance of secondary nucleation, and previous findings ^7,33,39^. Notably, there was no indication of larger preformed aggregates attaching to the fibril surface during elongation or secondary nucleation, implying that elongation occurs by attachment of single monomers.

### On the role of the flanking domains in secondary nucleation

The prevalence of secondary nucleation and branching shifted our attention to the nature of the interaction between free monomers and the fibril surface. An interesting aspect of HttEx1’s aggregation behaviour is that it is not only dependent on the length of the polyQ domain, but also the flanking segments Htt^NT^ and PRD. Htt^NT^ is generally expected to play an important driving force in the aggregation process, dramatically shortening the lag phase of aggregation (at a given protein concentration) ^24,26,27,55^. This implies a direct involvement in primary and/or secondary nucleation, consistent with a report that Htt^NT^-truncated HttEx1 constructs show a lack of lateral associations and branching ^31^. The role of Htt^NT^ can be studied by its removal prior to aggregation ^33^ yielding the construct ΔN15-HttEx1 (see Fig. 4A). The truncated HttEx1 aggregates slower than regular HttEx1 (see Fig. 4B), consistent with prior studies ^26,27,33,56,57^, resulting in a curve that is more similar to that of Q32-HttEx1 than Q44-HttEx1. Next, HS-AFM was used to analyse the fibril growth by the truncated ΔN15-Q44-HttEx1 by HS-AFM (see Fig. 4C-D). Interestingly, secondary nucleation followed by branching was also observed in these samples. Bi-directional elongation of the fibrillar structures was observed with an average rate of 5 ± 1 nm min^-1^, whereas secondary nucleated ΔN15-Q44-HttEx1 fibrils elongated with an average rate of 6 ± 1 nm min^-1^ corresponding well to the growth rates of the normal Q44-HttEx1 fibrils. A specific example of nucleated branching with an accompanying schematic is depicted in Figure 4D. Initially, small aggregates are visible on the fibril surface (green segments in Fig. 4D). These aggregates slowly grow in size to eventually take up a fibrillar structure. After elongating along the fibril, the growing end detaches from the parent-fibril, resulting in a branched structure. Another secondary nucleation site appears on top of the newly grown segment and shows the same phases of growth (red segment in 4D). The process of nucleated branching occurs with a similar frequency as in Q44-HttEx1 experiments. Lateral association of fibrils and growth on the surface of existing fibrils is therefore not solely the result of Htt^NT^ mediated association.

**Figure 4:**
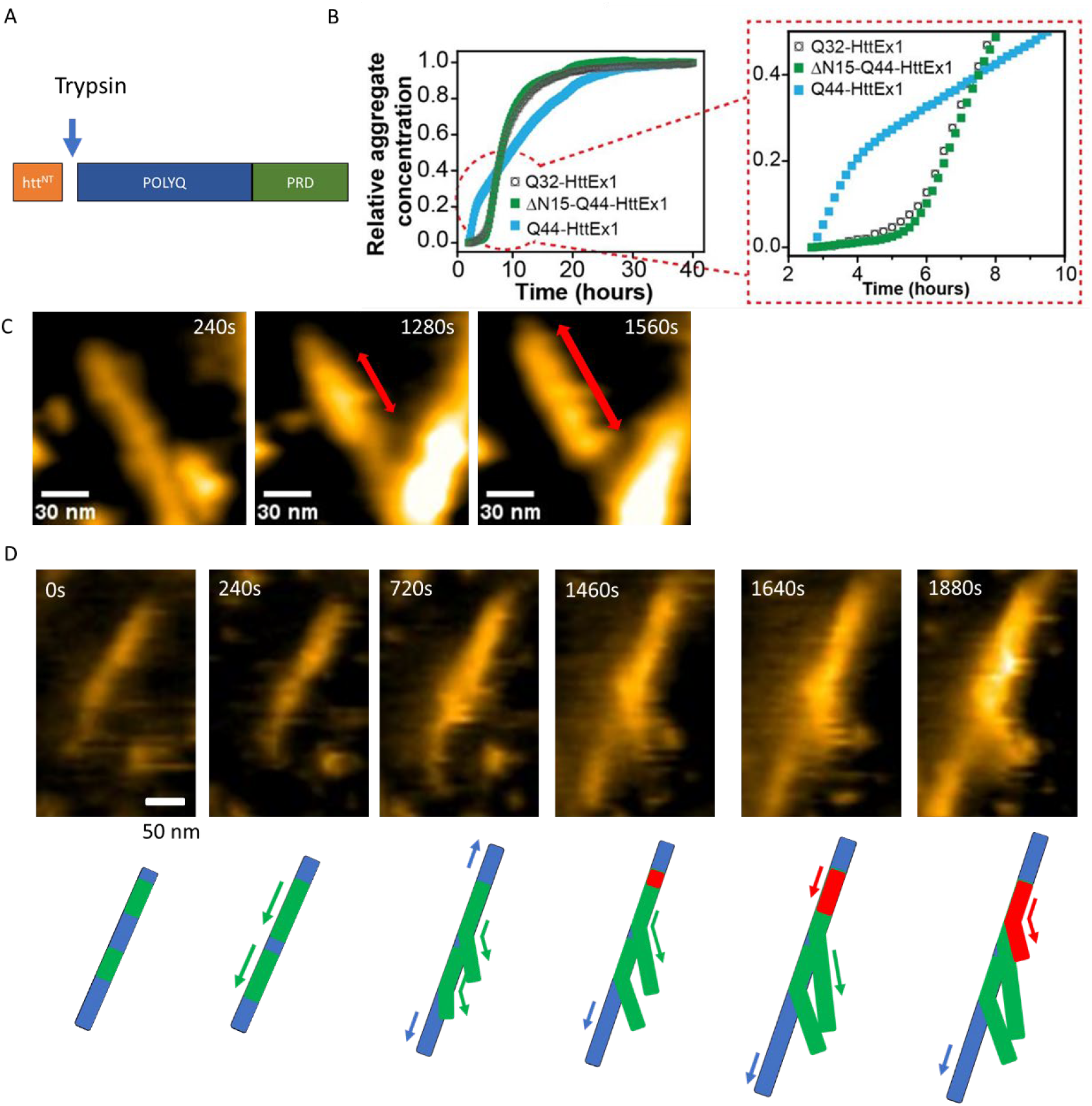
Real-time dynamics of seeded ΔN15-Q44-HttEx1 amyloid formation. A) Schematic ΔN15 Structure showing the cleavage site of Trypsin. B) Comparative aggregation kinetics of Q32-, Q44- and ΔN15-Q44-HttEx1, at 60 μM concentration, along with enlarged view of the initial lag and growth phase. All ThT data are measured in triplicates at 29°C and the mean of the triplicates is plotted. C) Snapshots of HS-AFM experiments showing secondary nucleation on a Δ15-Q44-HttEx1 amyloid fibril. D) Snapshots with corresponding schematic of secondary nucleation and subsequent branching on a Δ15-Q44-HttEx1 amyloid fibril. The blue fibril represents the parent fibrils, green fibrils are secondary nucleated fibrils and the red fibril is nucleated on top of the newly formed green fibrils. Scale bar 50 nm.

## Discussion

The notable role of secondary nucleation in the aggregation kinetics of HttEx1 was finally confirmed and visualised at the single fibril level in real-time. These results are in line with prior bulk and static studies on HttEx1 aggregation ^7,33^. We observed the complementary effects of polyQ length and flanking domains, in the complex aggregation pathway of these proteins ^27,57^. Morphological analysis by EM at various time intervals unveiled the gradual growth of amyloid-like fibrils, following initial formation of pre-fibrillar globular species, with fibrils becoming the dominant species by 20 hours. The globular species were attributed to pre-fibrillar oligomers described in earlier studies ^51,52^. Analysis of the fibril concentration via ThT-assay enabled the dissection of the fibril formation mechanism, which pointed to the prominence of secondary nucleation in this process. Next we probed the molecular underpinnings of the huntingtin amyloid formation in real time by HS-AFM, revealing the elongation and secondary nucleation pathways on the single-particle level. Consistent with bulk kinetics analyses, secondary nucleation of HttEx1 fibril formation was observed to occur on the sides of existing amyloid fibrils, being more common than direct primary nucleation. Thus, once fibrils are present (past the primary nucleation phase), this surface-mediated process appears to be a main driver of new fibril formation. Notably, this provides a direct observation of a molecular mechanism behind the concept of secondary nucleation: that the kinetics of fibril growth show a dependence on the total fibril mass, not just the number of growing ends. In our conventional understanding of (seeded) amyloid growth, the fibril ends would be considered the main (or only) source of fibril growth (aside from spontaneous primary nucleation events). The HS-AFM results highlight the importance of the available fibril surface area. The more fibrils that are already present, the more fibril surface area is available to capture the free available monomers, which will then misfold into the amyloid structure.

Due to these processes, small and isolated Q44-HttEx1 seeds were observed to grow into large bundles of fibrils as a result of monomer addition and subsequent branching. The appearance of large bundles due to secondary nucleation corresponds well to predictions from previous studies ^7^, where large bundles of fibrils were speculated to be a result of secondary nucleation rather than lateral association of existing fibrils. However, the driving force of branching remains uncertain. Nucleated branching has been speculated to result from a buildup of strain in the newly growing segment or steric constraints along the fibrils surface. Notably, the fibrils never completely separated in the analysed HS-AFM data. Thus, the different filaments remain attached together, forming higher-order structures reminiscent of fibril clusters seen in vitro ^36^ as well as in cells ^58^. Interestingly, this could imply that a single primary nucleation event could be behind the formation of entire μm-sized intracellular pathological inclusion bodies ^59^. Figure 5 provides a schematic visualization of this process ^32^.

**Figure 5.**
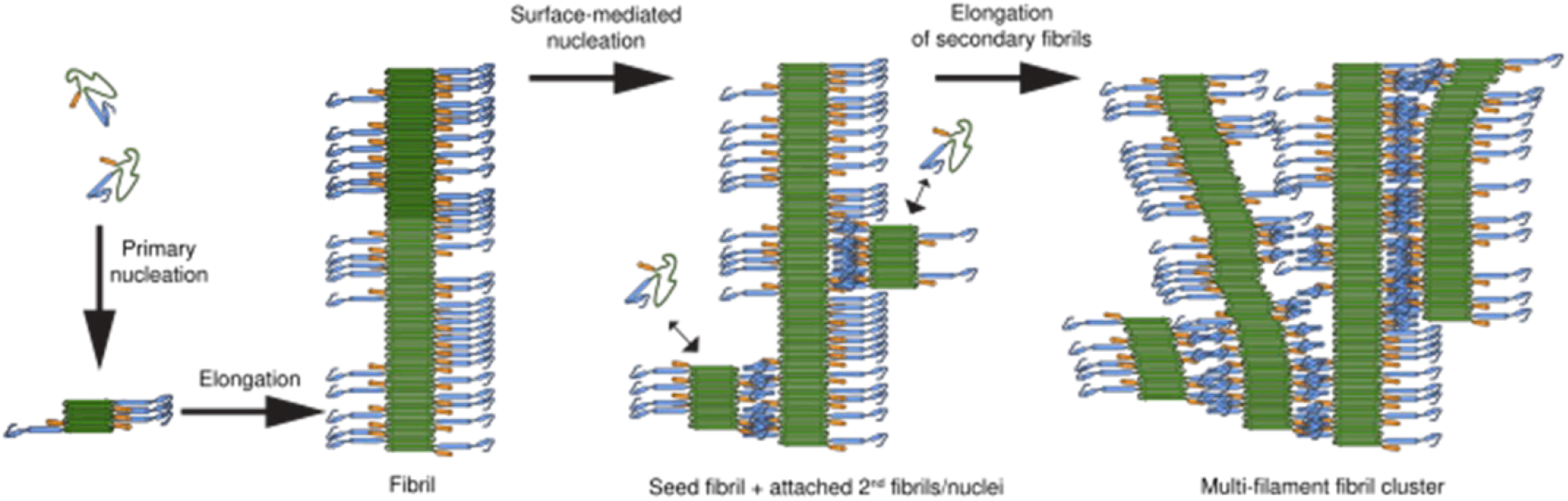
Schematic model for fibril assembly and secondary nucleation. Adapted from ref. ^33^ with permission.

Our results show that secondary nucleation happens along the fibril surface with no apparent specific sites, since multiple nucleation events can be observed on a single fibril, which all subsequently grow. The observed nucleation sites did not diffuse along the fibril end, which may indicate a reasonably strong adhesion between the seed fibril and the newly formed assembly. It is worth noting that we cannot exclude the possibility of monomers diffusing over the fibril surface ^60^, and nucleation resulting when the local concentration increases. The nucleation sites appeared gradually on the fibril surface, indicating that for Q44-HttEx1 the nucleation does not occur by association of large oligomers formed in solution to the fibrils. Instead, the nucleation of monomers into oligomers and finally the structural conversion to amyloidogenic form all seem to happen along the fibril surface for secondary nuclei.

Both fibril elongation as well as secondary nucleation and branching is still prevalent under HttEx1 deprived of Htt^NT^. One implication seems to be that the known aggregation-enhancing role of this flanking segment is not (primarily) derived from its role in secondary nucleation. This fits with prior models ^24,27,55^, which attribute the effect of Htt^NT^ to derive primarily from its effect on early stages of spontaneous aggregation (i.e., related to primary nucleation rather than secondary nucleation). Nonetheless, Htt^NT^ might still play a role in the surface reactivity of HttEx1 fibrils, yet it is not the solely responsible trigger for secondary nucleation. Our findings are in contrast with the results from Shen *et al*. ^31^, who reported that removing Htt^NT^ results in a lack of fibril bundling. On the contrary, Vieweg *et al*. ^26^ subsequently reported that removing Htt^NT^ induced a significant increase in lateral association and clumping of HttEx1 fibrils. Differences between previous studies may result from the different constructs used in preparation of the fibrils. Polymorphism of HttEx1 fibrils is known to be sensitive to the preparation conditions, differences in flanking segments and attached tags ^61^.

An important question relates to the part(s) of the amyloid structure that are responsible for the secondary nucleation on the surface. Here, it is important to revisit the architecture of HttEx1 fibrils, which feature a solid polyQ fibril core decorated on two sides by exposed flexible flanking segments, akin to a two-sided hairbrush design^28,33^ (Fig. 1B). Thus, there are two main types of surface features of HttEx1 fibrils that may be relevant. First, the water-facing surface of the block- or slab-like polyQ core itself. This core is multiple nm wide and assembled from multiple stacks of β-sheets, with at its surface the characteristic ‘grooved’ surface typical of amyloid fibrils. It is this grooved surface that is the binding site for ThT and similar ‘amyloid’ dyes. Alternatively, the flanking segments form a disordered and dynamic ‘fuzzy coat’ on two sides of this rigid core. This fuzzy coat is formed by the proline-rich C-terminus of HttEx1 and the N-terminal Htt^NT^ segment. In prior work, especially the PRD has been implicated in mediating interactions between protofilaments in HttEx1 fibril bundles ^33,53,62^. An interesting finding in the HS-AFM results was that the secondary filaments remain attached to the seed fibrils, which differs from prior studies of Ab42 aggregation where secondary protrusions were absent at the end of the reaction ^8^. The resulting fibril attachment is reminiscent of the side-by-side inter-filament interactions noted in prior work ^33^. This suggests a role for flanking domain interactions in this interaction, which may be present from the initial stages of the secondary nucleation process. In this context, the experiments with the N-terminally truncated proteins are notable, as they argue against a dominant role for the Htt^NT^. This may be considered surprising, since especially Htt^NT^ is widely implicated in the early stages of HttEx1 aggregation and nucleation ^24,27,55^. Instead, this may point to a role for the proline rich C-terminal domain. This may make sense, since this segment forms the bulk of the fibrillar fuzzy coat (being significantly larger than Htt^NT 28^). However, it is also known to prevent or reduce (spontaneous) aggregation from HttEx1 monomers ^24,63,64^. Further HS-AFM experiments with HttEx1 with removed or modified PRD domains could help elucidate the role of this C-terminal domain in the aggregation process and in particular secondary nucleation.

Our results demonstrate the importance of both elongation of the fibril ends and secondary nucleation along the fibril surface in HttEx1 amyloid growth. The prevalence of secondary nucleation and subsequent branching suggests that the majority of volume is created through secondary nucleation. This finding is consistent with mechanistic analysis of aggregation kinetics, but also provides a unique molecular and real-time perspective of the process. These results reinforce the idea that aggregation inhibitor studies should focus on prevention of secondary nucleation, as well as fibril elongation, in order to maximize therapeutic potential. Whilst largely unexplored, to our knowledge, creating compounds that could coat or pacify those parts of the fibril surface involved in secondary nucleation could be very promising for interfering with protein aggregation and prion-like propagation processes. This would be the case for HttEx1, as studied here, but may be extendable to other disease-relevant amyloids that are prone to secondary nucleation.

## Materials and Methods

### Protein expression and purification

A fusion protein construct having maltose-binding protein (MBP) as a solubility tag was used to produce Q32- and Q44-HttEx1 monomers, as previously described ^33,53^. The MBP-Q32-HttEx1, having 32 consecutive glutamines in polyQ domain was subcloned into a pMALc5x plasmid, whereas MBP-Q44-HttEx1, with 44 consecutive glutamines in polyQ domain was sub-cloned into pMALc2x plasmid by Genscript (Piscataway, NJ)[28, 51]. These constructs were expressed in Escherichia coli BL21(DE3) pLysS cells (Invitrogen, Grand Island, NY) and the cells were grown in LB medium with ampicillin and chloramphenicol at 37°C. The protein expression was induced by adding 0.6mM – 0.8mM of IPTG (Sigma) to the cell culture and shaking at 18°C for 16 hours. The cells were collected by pelleting them at 5250xg for 20mins. The cell lysis was done by suspending the cells in cold Phosphate Buffer Saline (PBS), pH 7.4 buffer with 0.5mg/ml of lysozyme, 0.01% sodium azide and 1 Pierce EDTA-free protease inhibitor tablet (ThermoFisher Scientific). This suspension was then sonicated to burst the cells using VCX130 Vibra-cell sonicator, Sonics & Materials, Inc by applying 70% amplitude for 20mins (pulse alternation – 10s pulse, 10s break). The cell debris was removed by centrifuging the lysate at 125,000xg for 45 mins. The supernatant containing soluble fusion protein was collected, filtered with 0.45μm syringe filters and purified using a HisTrap HP nickel column (GE Healthcare) on an Akta FPLC system. The protein was eluted using an imidazole gradient (from 10 to 75%) and the protein presence and purity was verified by 12% SDS-gel as described in ^33^. The purified fused protein was collected and imidazole was removed using dialysis (Dialysis tubing, MWCO 12.4kDa, Sigma-Aldrich). To determine the protein concentration, the absorbance was measured at 280nm with a JASCO V-750 spectrophotometer. The absorbance was measured in triplicate and average of the absorbance values (n=3) was taken and divided by the extinction coefficient of 66,350 M^-1^cm^-1^ (derived from the fusion protein sequence by ProtParam tool by ExPASy ^65^). The molecular weight was verified by ESI-TOF mass spectrometry ^33^.

### Protein aggregation

MBP-Q32-HttEx1 and MBP-Q44-HttEx1 were cleaved to release soluble HttEx1 by using Factor Xa (FXa; Promega, Madison, WI) at 400:1 (Fused protein: Protease) molar ratio. To generate ΔN15-Q44-HttEx1, the MBP-Q44-HttEx1 fusion protein was cleaved by Trypsin (N-tosyl-L-phenylalanine chloromethyl ketone treated trypsin lyophilized powder from bovine pancreas), re-dissolved in PBS buffer with 3μM HCl) at 50:1 (Fused protein: Protease) molar ratio. The protein cleavage was carried out at room temperature and was monitored by SDS-PAGE (Bio-Rad Mini-Protein Precast TGX Gels 12%). The final protein aggregates were collected after 4 days, and pelleted down by centrifuging at 2400xg for 20 mins. The supernatant was removed and the protein aggregates were washed three times with PBS buffer to ensure complete removal of MBP. The pre-existing Q44-HttEx1 fibril seeds used in AFM and kinetics measurements were prepared by aggregating 50μM MBP-Q44-HttEx1 fused protein with FXa protease at 400:1 (Fused protein: Protease) molar ratio. The aggregation was performed at room temperature for 96 hours. After that, the protein aggregates were washed three times using PBS buffer.

### High-Speed Atomic Force Microscopy

Atomic force microscopy was performed using an SS-NEX High-Speed Atomic Force Microscope (RIBM, Japan) in amplitude modulation tapping mode in liquid. Ultra-short cantilevers with high resonance frequency (USC-F1.2-k0.15, Nanoworld, Switzerland) were used. Short Q44-HttEx1 fibrils (seeds), which were preformed at room temperature, were incubated on mica surface for 10 minutes, and subsequently washed with PBS. The sample is then placed in the recording chamber containing the cantilever in a solution containing 60μM MBP-Q44-HttEx1 cleaved with either FXa at 400:1 molar ratio (HttEx1:FXa), or trypsin 50:1 (HttEx1:Trypsin) molar ratio. Images were recorded every 5 seconds, unless stated otherwise. All images were processed using ImageJ ^66^ software with additional home-written plugins.

### Thioflavin T (ThT) Assay

The aggregation kinetics were measured using 20μM, 30μM, 50μM and 60μM MBP-Q32-HttEx1, with cleavage by FXa at 400:1 molar ratio (HttEx1: FXa). These ThT assays were performed with the ThT present during the aggregation reaction, using fluorescence plate readers, as described next. For kinetics assays, the 60μM MBP-Q44-HttEx1 was divided in two batches: the first batch was treated with FXa at 400:1 molar ratio (HttEx1: FXa) to study the Q44-HttEx1 aggregation kinetics, and the other batch was treated with Trypsin (N-tosyl-L-phenylalanine chloromethyl ketone treated trypsin lyophilized powder from bovine pancreas, re-dissolved in PBS buffer with 3μM HCl) at 50:1 molar ratio (HttEx1: trypsin) to study the ΔN15-Q44-HttEx1 aggregation kinetics. The effects of seeds on aggregation kinetics of 67.5μM fused Q44-HttEx1 were measured by cleaving it by 400:1 molar ratio (HttEx1: FXa), adding seeds of preformed Q44-HttEx1 fibrils made at room temperature. The seed concentration used for the assay was 10%. A 1mM ThT stock solution was made in DMSO and the final concentration of ThT in the well was 15μM. The experiments were carried out in black polystyrene, clear bottom, 96-well plates (Corning, U.S.A) and measured on a TECAN Spark 10M microplate reader. The excitation wavelength is 442nm and the emission wavelength is 484nm, with excitation and emission bandwidths of 10nm. The experiments were done in triplicate.

### Negative-stain TEM

Negative-stain TEM was performed on samples from the Q32-HttEx1 aggregation process, at respective time intervals (2hours, 20hours, 71hours and 96hours) (Figure S1C), as well as the final Q44-HttEx1 (Figure 2C), and seeded and unseeded Q44-HttEx1 fibrils (Figure S2, B&C). The time dependent samples were frozen on specific time points in liquid nitrogen, and thawed at room temperature for making grids. The moment the sample was thawed, it was quickly mixed and placed on the grid. The plain carbon support film – 200 mesh copper grids (SKU FCF200-Cu-50, Electron Microscopy Sciences, Hatfield, PA) were glow discharged for 0.5 - 1min and then the fibril samples were deposited on the grid for 0.5 – 1min. The excess sample was removed by blotting and then the negative staining agent 2% (w/v) uranyl acetate was allowed to stain the grid for 0.5 – 1min. The excess stain was removed by blotting and the grid was air dried. The grid was imaged on Philips CM120 electron microscope operating at 120kV with a slow scan CCD camera (Gatan). Fiji ^67^, was used to measure the fibril widths, analogous to prior work^33^. Each measurement was transverse to the fibril long-axis, and was the measure of the negative stained area of the fibril. In TEM images shown, the fibrils widths were determined in the regions where the fibril boundaries are clearly defined. The fibril widths were determined on isolated fibrils, using Plot profile tool in Fiji which plots the average grayscale intensity. We obtained three measurements per fibril, with exception to the fibrils where the width varied significantly. For the fibrils that varied in width, we did more than three measurements per fibril.

## Acknowledgments

W.H.R. is supported by a Dieptestrategie grant of the Zernike Institute National Research Centre of the Rijksuniversiteit Groningen and by the EU funded project MOSBRI (101004806). G.J. and P.C.A.v.d.W. acknowledge the financial support from the CampagneTeam Huntington Foundation. We thank Marc Stuart for help with the use of the EM facility and Gea Schuurman-Wolters for her help with the biochemistry facility instruments.

## References

(1) Soto, C. Unfolding the Role of Protein Misfolding in Neurodegenerative Diseases. Nat Rev Neurosci 2003, 4 (1), 49–60. 10.1038/nrn1007.

(2) Törnquist, M.; Michaels, T. C. T.; Sanagavarapu, K.; Yang, X.; Meisl, G.; Cohen, S. I. A.; Knowles, T. P. J.; Linse, S. Secondary Nucleation in Amyloid Formation. Chemical Communications 2018, 54 (63), 8667–8684. 10.1039/C8CC02204F.

(3) Sinnige, T. Molecular Mechanisms of Amyloid Formation in Living Systems. Chem Sci 2022, 13 (24), 7080–7097. 10.1039/D2SC01278B.

(4) Kakkar, V.; Månsson, C.; de Mattos, E. P.; Bergink, S.; van der Zwaag, M.; van Waarde, M. A. W. H.; Kloosterhuis, N. J.; Melki, R.; van Cruchten, R. T. P.; Al-Karadaghi, S.; Arosio, P.; Dobson, C. M.; Knowles, T. P. J.; Bates, G. P.; van Deursen, J. M.; Linse, S.; van de Sluis, B.; Emanuelsson, C.; Kampinga, H. H. The S/T-Rich Motif in the DNAJB6 Chaperone Delays Polyglutamine Aggregation and the Onset of Disease in a Mouse Model. Mol Cell 2016, 62 (2), 272–283. 10.1016/j.molcel.2016.03.017.

(5) Rodriguez Camargo, D. C.; Sileikis, E.; Chia, S.; Axell, E.; Bernfur, K.; Cataldi, R. L.; Cohen, S. I. A.; Meisl, G.; Habchi, J.; Knowles, T. P. J.; Vendruscolo, M.; Linse, S. Proliferation of Tau 304–380 Fragment Aggregates through Autocatalytic Secondary Nucleation. ACS Chem Neurosci 2021, 12 (23), 4406–4415. 10.1021/acschemneuro.1c00454.

(6) Cohen, S. I. A.; Linse, S.; Luheshi, L. M.; Hellstrand, E.; White, D. A.; Rajah, L.; Otzen, D. E.; Vendruscolo, M.; Dobson, C. M.; Knowles, T. P. J. Proliferation of Amyloid-Β42 Aggregates Occurs through a Secondary Nucleation Mechanism. Proceedings of the National Academy of Sciences 2013, 110 (24), 9758–9763. 10.1073/pnas.1218402110.

(7) Wagner, A. S.; Politi, A. Z.; Ast, A.; Bravo-rodriguez, K.; Baum, K.; Buntru, A.; Strempel, N. U.; Brusendorf, L.; Hänig, C.; Boeddrich, A.; Plassmann, S.; Klockmeier, K.; Ramirez-anguita, J. M.; Sanchez-garcia, E.; Wolf, J.; Wanker, E. E. Self-Assembly of Mutant Huntingtin Exon-1 Fragments into Large Complex Fibrillar Structures Involves Nucleated Branching. J Mol Biol 2018, 430 (12), 1725–1744. 10.1016/j.jmb.2018.03.017.

(8) Törnquist, M.; Cukalevski, R.; Weininger, U.; Meisl, G.; Knowles, T. P. J.; Leiding, T.; Malmendal, A.; Akke, M.; Linse, S. Ultrastructural Evidence for Self-Replication of Alzheimer-Associated Aβ42 Amyloid along the Sides of Fibrils. Proceedings of the National Academy of Sciences 2020, 117 (21), 11265–11273. 10.1073/pnas.1918481117.

(9) Hadi Alijanvand, S.; Peduzzo, A.; Buell, A. K. Secondary Nucleation and the Conservation of Structural Characteristics of Amyloid Fibril Strains. Front Mol Biosci 2021, 8:669994. 10.3389/fmolb.2021.669994.

(10) Staats, R.; Brotzakis, Z. F.; Chia, S.; Horne, R. I.; Vendruscolo, M. Optimization of a Small Molecule Inhibitor of Secondary Nucleation in α-Synuclein Aggregation. Front Mol Biosci 2023, 10:1155753. 10.3389/fmolb.2023.1155753.

(11) Michaels, T. C. T.; Dear, A. J.; Cohen, S. I. A.; Vendruscolo, M.; Knowles, T. P. J. Kinetic Profiling of Therapeutic Strategies for Inhibiting the Formation of Amyloid Oligomers. J Chem Phys 2022, 156 (16), 164904. 10.1063/5.0077609.

(12) Michaels, T. C. T.; Šarić, A.; Meisl, G.; Heller, G. T.; Curk, S.; Arosio, P.; Linse, S.; Dobson, C. M.; Vendruscolo, M.; Knowles, T. P. J. Thermodynamic and Kinetic Design Principles for Amyloid-Aggregation Inhibitors. Proceedings of the National Academy of Sciences 2020, 117 (39), 24251–24257. 10.1073/pnas.2006684117.

(13) Doig, A. J.; del Castillo-Frias, M. P.; Berthoumieu, O.; Tarus, B.; Nasica-Labouze, J.; Sterpone, F.; Nguyen, P. H.; Hooper, N. M.; Faller, P.; Derreumaux, P. Why Is Research on Amyloid-β Failing to Give New Drugs for Alzheimer’s Disease? ACS Chem Neurosci 2017, 8 (7), 1435–1437. 10.1021/acschemneuro.7b00188.

(14) Bates, G. P.; Dorsey, R.; Gusella, J. F.; Hayden, M. R.; Kay, C.; Leavitt, B. R.; Nance, M.; Ross, C. A.; Scahill, R. I.; Wetzel, R.; Wild, E. J.; Tabrizi, S. J. Huntington Disease. Nat Rev Dis Primers 2015, 1 (1), 15005. 10.1038/nrdp.2015.5.

(15) DiFiglia, M.; Sapp, E.; Chase, K. O.; Davies, S. W.; Bates, G. P.; Vonsattel, J. P.; Aronin, N. Aggregation of Huntingtin in Neuronal Intranuclear Inclusions and Dystrophic Neurites in Brain. Science (1979) 1997, 277 (5334), 1990–1993. 10.1126/science.277.5334.1990.

(16) Arrasate, M.; Mitra, S.; Schweitzer, E. S.; Segal, M. R.; Finkbeiner, S. Inclusion Body Formation Reduces Levels of Mutant Huntingtin and the Risk of Neuronal Death. Nature 2004, 431 (7010), 805–810. 10.1038/nature02998.

(17) Drombosky, K. W.; Rode, S.; Kodali, R.; Jacob, T. C.; Palladino, M. J.; Wetzel, R. Mutational Analysis Implicates the Amyloid Fibril as the Toxic Entity in Huntington’s Disease. Neurobiol Dis 2018, 120, 126–138. 10.1016/j.nbd.2018.08.019.

(18) Ruggeri, F. S.; Vieweg, S.; Cendrowska, U.; Longo, G.; Chiki, A.; Lashuel, H. A.; Dietler, G. Nanoscale Studies Link Amyloid Maturity with Polyglutamine Diseases Onset. Sci Rep 2016, 6 (1), 31155. 10.1038/srep31155.

(19) Neueder, A.; Landles, C.; Ghosh, R.; Howland, D.; Myers, R. H.; Faull, R. L. M.; Tabrizi, S. J.; Bates, G. P. The Pathogenic Exon 1 HTT Protein Is Produced by Incomplete Splicing in Huntington’s Disease Patients. Sci Rep 2017, 7 (1), 1307. 10.1038/s41598-017-01510-z.

(20) Sathasivam, K.; Neueder, A.; Gipson, T. A.; Landles, C.; Benjamin, A. C.; Bondulich, M. K.; Smith, D. L.; Faull, R. L. M.; Roos, R. A. C.; Howland, D.; Detloff, P. J.; Housman, D. E.; Bates, G. P. Aberrant Splicing of HTT Generates the Pathogenic Exon 1 Protein in Huntington Disease. Proceedings of the National Academy of Sciences 2013, 110 (6), 2366–2370. 10.1073/pnas.1221891110.

(21) Landles, C.; Sathasivam, K.; Weiss, A.; Woodman, B.; Moffitt, H.; Finkbeiner, S.; Sun, B.; Gafni, J.; Ellerby, L. M.; Trottier, Y.; Richards, W. G.; Osmand, A.; Paganetti, P.; Bates, G. P. Proteolysis of Mutant Huntingtin Produces an Exon 1 Fragment That Accumulates as an Aggregated Protein in Neuronal Nuclei in Huntington Disease. Journal of Biological Chemistry 2010, 285 (12), 8808–8823. 10.1074/jbc.M109.075028.

(22) Kennedy, L.; Evans, E.; Chen, C.-M.; Craven, L.; Detloff, P. J.; Ennis, M.; Shelbourne, P. F. Dramatic Tissue-Specific Mutation Length Increases Are an Early Molecular Event in Huntington Disease Pathogenesis. Hum Mol Genet 2003, 12 (24), 3359–3367. 10.1093/hmg/ddg352.

(23) Lin, H.-K.; Boatz, J. C.; Krabbendam, I. E.; Kodali, R.; Hou, Z.; Wetzel, R.; Dolga, A. M.; Poirier, M. A.; van der Wel, P. C. A. Fibril Polymorphism Affects Immobilized Non-Amyloid Flanking Domains of Huntingtin Exon1 Rather than Its Polyglutamine Core. Nat Commun 2017, 8 (1), 15462. 10.1038/ncomms15462.

(24) Crick, S. L.; Ruff, K. M.; Garai, K.; Frieden, C.; Pappu, R. V. Unmasking the Roles of N- and C-Terminal Flanking Sequences from Exon 1 of Huntingtin as Modulators of Polyglutamine Aggregation. Proceedings of the National Academy of Sciences 2013, 110 (50), 20075–20080. 10.1073/pnas.1320626110.

(25) Arndt, J. R.; Chaibva, M.; Legleiter, J. The Emerging Role of the First 17 Amino Acids of Huntingtin in Huntington’s Disease. Biomol Concepts 2015, 6 (1), 33–46. 10.1515/bmc-2015-0001.

(26) Vieweg, S.; Mahul-Mellier, A. L.; Ruggeri, F. S.; Riguet, N.; DeGuire, S. M.; Chiki, A.; Cendrowska, U.; Dietler, G.; Lashuel, H. A. The Nt17 Domain and Its Helical Conformation Regulate the Aggregation, Cellular Properties and Neurotoxicity of Mutant Huntingtin Exon 1. J Mol Biol 2021, 433 (21), 167222. 10.1016/j.jmb.2021.167222.

(27) Thakur, A. K.; Jayaraman, M.; Mishra, R.; Thakur, M.; Chellgren, V. M.; L Byeon, I.-J.; Anjum, D. H.; Kodali, R.; Creamer, T. P.; Conway, J. F.; M Gronenborn, A.; Wetzel, R. Polyglutamine Disruption of the Huntingtin Exon 1 N Terminus Triggers a Complex Aggregation Mechanism. Nat Struct Mol Biol 2009, 16 (4), 380–389. 10.1038/nsmb.1570.

(28) Bagherpoor Helabad, M.; Matlahov, I.; Kumar, R.; Daldrop, J. O.; Jain, G.; Weingarth, M.; van der Wel, P. C. A.; Miettinen, M. S. Integrative Determination of the Atomic Structure of Mutant Huntingtin Exon 1 Fibrils Implicated in Huntington’s Disease. bioRxiv 2024, 2023.07.21.549993. 10.1101/2023.07.21.549993.

(29) Ceccon, A.; Tugarinov, V.; Torricella, F.; Clore, G. M. Quantitative NMR Analysis of the Kinetics of Prenucleation Oligomerization and Aggregation of Pathogenic Huntingtin Exon-1 Protein. Proc Natl Acad Sci U S A 2022, 119 (29), e2207690119. 10.1073/pnas.2207690119.

(30) Tam, S.; Spiess, C.; Auyeung, W.; Joachimiak, L.; Chen, B.; Poirier, M. A.; Frydman, J. The Chaperonin TRiC Blocks a Huntingtin Sequence Element That Promotes the Conformational Switch to Aggregation. Nat Struct Mol Biol 2009, 16 (12), 1279–1285. 10.1038/nsmb.1700.

(31) Shen, K.; Calamini, B.; Fauerbach, J. A.; Ma, B.; Shahmoradian, S. H.; Serrano Lachapel, I. L.; Chiu, W.; Lo, D. C.; Frydman, J. Control of the Structural Landscape and Neuronal Proteotoxicity of Mutant Huntingtin by Domains Flanking the PolyQ Tract. Elife 2016, 5, e18065. 10.7554/eLife.18065.

(32) Gottlieb, L.; Guo, L.; Shorter, J.; Marmorstein, R. N-Alpha-Acetylation of Huntingtin Protein Increases Its Propensity to Aggregate. Journal of Biological Chemistry 2021, 297 (6), 101363. 10.1016/j.jbc.2021.101363.

(33) Boatz, J. C.; Piretra, T.; Lasorsa, A.; Matlahov, I.; Conway, J. F.; Wel, P. C. A. Van Der. Protofilament Structure and Supramolecular Polymorphism of Aggregated Mutant Huntingtin Exon 1. J Mol Biol 2020, 432 (16), 4722–4744. 10.1016/j.jmb.2020.06.021.

(34) Nazarov, S.; Chiki, A.; Boudeffa, D.; Lashuel, H. A. Structural Basis of Huntingtin Fibril Polymorphism Revealed by Cryogenic Electron Microscopy of Exon 1 HTT Fibrils. J Am Chem Soc 2022, 144 (24), 10723–10735. 10.1021/jacs.2c00509.

(35) Isas, J. M.; Pandey, N. K.; Xu, H.; Teranishi, K.; Okada, A. K.; Fultz, E. K.; Rawat, A.; Applebaum, A.; Meier, F.; Chen, J.; Langen, R.; Siemer, A. B. Huntingtin Fibrils with Different Toxicity, Structure, and Seeding Potential Can Be Interconverted. Nat Commun No. 2021, 1–11. 10.1038/s41467-021-24411-2.

(36) Galaz-Montoya, J. G.; Shahmoradian, S. H.; Shen, K.; Frydman, J.; Chiu, W. Cryo-Electron Tomography Provides Topological Insights into Mutant Huntingtin Exon 1 and PolyQ Aggregates. Commun Biol 2021, 4 (1), 849. 10.1038/s42003-021-02360-2.

(37) Nekooki-Machida, Y.; Kurosawa, M.; Nukina, N.; Ito, K.; Oda, T.; Tanaka, M. Distinct Conformations of in Vitro and in Vivo Amyloids of Huntingtin-Exon1 Show Different Cytotoxicity. Proceedings of the National Academy of Sciences 2009, 106 (24), 9679–9684. 10.1073/pnas.0812083106.

(38) Mario Isas, J.; Pandey, N. K.; Xu, H.; Teranishi, K.; Okada, A. K.; Fultz, E. K.; Rawat, A.; Applebaum, A.; Meier, F.; Chen, J.; Langen, R.; Siemer, A. B. Huntingtin Fibrils with Different Toxicity, Structure, and Seeding Potential Can Be Interconverted. Nat Commun 2021, 12 (1), 4272. 10.1038/s41467-021-24411-2.

(39) Ast, A.; Buntru, A.; Schindler, F.; Hasenkopf, R.; Schulz, A.; Brusendorf, L.; Klockmeier, K.; Grelle, G.; McMahon, B.; Niederlechner, H.; Jansen, I.; Diez, L.; Edel, J.; Boeddrich, A.; Franklin, S. A.; Baldo, B.; Schnoegl, S.; Kunz, S.; Purfürst, B.; Gaertner, A.; Kampinga, H. H.; Morton, A. J.; Petersén, Å.; Kirstein, J.; Bates, G. P.; Wanker, E. E. MHTT Seeding Activity: A Marker of Disease Progression and Neurotoxicity in Models of Huntington’s Disease. Mol Cell 2018, 71 (5), 675–688.e6. 10.1016/j.molcel.2018.07.032.

(40) Walker, L. C.; LeVine, H. Corruption and Spread of Pathogenic Proteins in Neurodegenerative Diseases*. Journal of Biological Chemistry 2012, 287 (40), 33109–33115. 10.1074/jbc.R112.399378.

(41) Pearce, M. M. P.; Kopito, R. R. Prion-like Characteristics of Polyglutamine-Containing Proteins. Cold Spring Harb Perspect Med 2018. 10.1101/cshperspect.a024257.

(42) Dahlgren, P. R.; Karymov, M. A.; Bankston, J.; Holden, T.; Thumfort, P.; Ingram, V. M.; Lyubchenko, Y. L. Atomic Force Microscopy Analysis of the Huntington Protein Nanofibril Formation. Nanomedicine 2005, 1 (1), 52–57. 10.1016/J.NANO.2004.11.004.

(43) Ando, T.; Uchihashi, T.; Scheuring, S. Filming Biomolecular Processes by High-Speed Atomic Force Microscopy. Chem Rev 2014, 114 (6), 3120–3188. 10.1021/cr4003837.

(44) van Ewijk, C.; Xu, F.; Maity, S.; Sheng, J.; Stuart, M. C. A.; Feringa, B. L.; Roos, W. H. Light-Triggered Disassembly of Molecular Motor-Based Supramolecular Polymers Revealed by High-Speed AFM. Angewandte Chemie International Edition 2024, 63 (14), e202319387. 10.1002/anie.202319387.

(45) Maity, S.; Trinco, G.; Buzón, P.; Anshari, Z. R.; Kodera, N.; Ngo, K. X.; Ando, T.; Slotboom, D. J.; Roos, W. H. High-Speed Atomic Force Microscopy Reveals a Three-State Elevator Mechanism in the Citrate Transporter CitS. Proceedings of the National Academy of Sciences 2022, 119 (6), e2113927119. 10.1073/pnas.2113927119.

(46) Watanabe-Nakayama, T.; Tsuji, M.; Umeda, K.; Oguchi, T.; Konno, H.; Noguchi-Shinohara, M.; Kiuchi, Y.; Kodera, N.; Teplow, D. B.; Ono, K. Structural Dynamics of Amyloid-β Protofibrils and Actions of Anti-Amyloid-β Antibodies as Observed by High-Speed Atomic Force Microscopy. Nano Lett 2023, 23 (13), 6259–6268. 10.1021/acs.nanolett.3c00187.

(47) Banerjee, S.; Sun, Z.; Hayden, E. Y.; Teplow, D. B.; Lyubchenko, Y. L. Nanoscale Dynamics of Amyloid β-42 Oligomers As Revealed by High-Speed Atomic Force Microscopy. ACS Nano 2017, 11 (12), 12202–12209. 10.1021/acsnano.7b05434.

(48) Milhiet, P.-E.; Yamamoto, D.; Berthoumieu, O.; Dosset, P.; Le Grimellec, C.; Verdier, J.-M.; Marchal, S.; Ando, T. Deciphering the Structure, Growth and Assembly of Amyloid-Like Fibrils Using High-Speed Atomic Force Microscopy. PLoS One 2010, 5 (10), e13240.

(49) Jain, G.; Trombetta-Lima, M.; Matlahov, I.; Ribas, H. T.; Dolga, A. M.; van der Wel, P. C. A. Inhibitor-Based Modulation of Huntingtin Aggregation Reduces Fibril Toxicity. bioRxiv 2023, 2023.04.24.537565. 10.1101/2023.04.24.537565.

(50) Meisl, G.; Kirkegaard, J. B.; Arosio, P.; Michaels, T. C. T.; Vendruscolo, M.; Dobson, C. M.; Linse, S.; Knowles, T. P. J. Molecular Mechanisms of Protein Aggregation from Global Fitting of Kinetic Models. Nat Protoc 2016, 11 (2), 252–272. 10.1038/nprot.2016.010.

(51) Nucifora, L. G.; Burke, K. A.; Feng, X.; Arbez, N.; Zhu, S.; Miller, J.; Yang, G.; Ratovitski, T.; Delannoy, M.; Muchowski, P. J.; Finkbeiner, S.; Legleiter, J.; Ross, C. A.; Poirier, M. A. Identification of Novel Potentially Toxic Oligomers Formed in Vitro from Mammalian-Derived Expanded Huntingtin Exon-1 Protein*. Journal of Biological Chemistry 2012, 287 (19), 16017–16028. 10.1074/jbc.M111.252577.

(52) Legleiter, J.; Mitchell, E.; Lotz, G. P.; Sapp, E.; Ng, C.; DiFiglia, M.; Thompson, L. M.; Muchowski, P. J. Mutant Huntingtin Fragments Form Oligomers in a Polyglutamine Length-Dependent Manner in Vitro and in Vivo*. Journal of Biological Chemistry 2010, 285 (19), 14777–14790. 10.1074/jbc.M109.093708.

(53) Lin, H. K.; Boatz, J. C.; Krabbendam, I. E.; Kodali, R.; Hou, Z.; Wetzel, R.; Dolga, A. M.; Poirier, M. A.; Van Der Wel, P. C. A. Fibril Polymorphism Affects Immobilized Non-Amyloid Flanking Domains of Huntingtin Exon1 Rather than Its Polyglutamine Core. Nat Commun 2017, 8 (May), 1–12. 10.1038/ncomms15462.

(54) Hoop, C. L.; Lin, H.-K.; Kar, K.; Magyarfalvi, G.; Lamley, J. M.; Boatz, J. C.; Mandal, A.; Lewandowski, J. R.; Wetzel, R.; van der Wel, P. C. A. Huntingtin Exon 1 Fibrils Feature an Interdigitated β-Hairpin–Based Polyglutamine Core. Proceedings of the National Academy of Sciences 2016, 113 (6), 1546–1551. 10.1073/pnas.1521933113.

(55) Torricella, F.; Tugarinov, V.; Clore, G. M. Nucleation of Huntingtin Aggregation Proceeds via Conformational Conversion of Pre-Formed, Sparsely-Populated Tetramers. Advanced Science 2024, 11 (24), 2309217. 10.1002/advs.202309217.

(56) Sivanandam, V. N.; Jayaraman, M.; Hoop, C. L.; Kodali, R.; Wetzel, R.; van der Wel, P. C. A. The Aggregation-Enhancing Huntingtin N-Terminus Is Helical in Amyloid Fibrils. J Am Chem Soc 2011, 133 (12), 4558–4566. 10.1021/ja110715f.

(57) Jayaraman, M.; Mishra, R.; Kodali, R.; Thakur, A. K.; Koharudin, L. M. I.; Gronenborn, A. M.; Wetzel, R. Kinetically Competing Huntingtin Aggregation Pathways Control Amyloid Polymorphism and Properties. Biochemistry 2012, 51 (13), 2706–2716. 10.1021/bi3000929.

(58) Peskett, T. R.; Rau, F.; O’Driscoll, J.; Patani, R.; Lowe, A. R.; Saibil, H. R. A Liquid to Solid Phase Transition Underlying Pathological Huntingtin Exon1 Aggregation. Mol Cell 2018, 70 (4), 588–601.e6. 10.1016/j.molcel.2018.04.007.

(59) Sinnige, T.; Meisl, G.; Michaels, T. C. T.; Vendruscolo, M.; Knowles, T. P. J.; Morimoto, R. I. Kinetic Analysis Reveals That Independent Nucleation Events Determine the Progression of Polyglutamine Aggregation in C. Elegans. Proceedings of the National Academy of Sciences 2021, 118 (11), e2021888118. 10.1073/pnas.2021888118.

(60) Maity, S.; Ottelé, J.; Santiago, G. M.; Frederix, P. W. J. M.; Kroon, P.; Markovitch, O.; Stuart, M. C. A.; Marrink, S. J.; Otto, S.; Roos, W. H. Caught in the Act: Mechanistic Insight into Supramolecular Polymerization-Driven Self-Replication from Real-Time Visualization. J Am Chem Soc 2020, 142 (32), 13709–13717. 10.1021/jacs.0c02635.

(61) Matlahov, I.; van der Wel, P. C. A. Conformational Studies of Pathogenic Expanded Polyglutamine Protein Deposits from Huntington’s Disease. Exp Biol Med 2019, 1, 1584–1595. 10.1177/1535370219856620.

(62) Bugg, C. W.; Isas, J. M.; Fischer, T.; Patterson, P. H.; Langen, R. Structural Features and Domain Organization of Huntingtin Fibrils. Journal of Biological Chemistry 2012, 287 (38), 31739–31746. 10.1074/jbc.M112.353839.

(63) Bhattacharyya, A.; Thakur, A. K.; Chellgren, V. M.; Thiagarajan, G.; Williams, A. D.; Chellgren, B. W.; Creamer, T. P.; Wetzel, R. Oligoproline Effects on Polyglutamine Conformation and Aggregation. J Mol Biol 2006, 355 (3), 524–535. 10.1016/j.jmb.2005.10.053.

(64) Darnell, G.; Orgel, J. P. R. O.; Pahl, R.; Meredith, S. C. Flanking Polyproline Sequences Inhibit β-Sheet Structure in Polyglutamine Segments by Inducing PPII-like Helix Structure. J Mol Biol 2007, 374 (3), 688–704. 10.1016/j.jmb.2007.09.023.

(65) Gasteiger, E.; Gattiker, A.; Hoogland, C.; Ivanyi, I.; Appel, R. D.; Bairoch, A. ExPASy: The Proteomics Server for in-Depth Protein Knowledge and Analysis. Nucleic Acids Res 2003, 31 (13), 3784–3788. 10.1093/nar/gkg563.

(66) Abràmoff, M. D.; Magalhães, P. J.; Ram, S. J. Image Processing with ImageJ. Biophotonics international 2004, 11 (7), 36–42.

(67) Schindelin, J.; Arganda-Carreras, I.; Frise, E.; Kaynig, V.; Longair, M.; Pietzsch, T.; Preibisch, S.; Rueden, C.; Saalfeld, S.; Schmid, B.; Tinevez, J.-Y.; White, D. J.; Hartenstein, V.; Eliceiri, K.; Tomancak, P.; Cardona, A. Fiji: An Open-Source Platform for Biological-Image Analysis. Nat Methods 2012, 9 (7), 676–682. 10.1038/nmeth.2019.

